# Persistent activation of interlinked Th2-airway epithelial gene networks in sputum-derived cells from aeroallergen-sensitized symptomatic atopic asthmatics

**DOI:** 10.1101/063602

**Authors:** Anya C. Jones, Niamh M. Troy, Elisha White, Elysia M. Hollams, Alexander M. Gout, Kak-Ming Ling, Anthony Kicic, Peter D Sly, Patrick G Holt, Graham L Hall, Anthony Bosco

## Abstract

**Rationale:** Atopic asthma is a persistent disease characterized by intermittent wheeze and progressive loss of lung function. The disease is thought to be driven primarily by chronic aeroallergen-induced Th2-associated airways inflammation. However, the vast majority of atopics do not develop asthma-related wheeze, despite ongoing exposure to aeroallergens to which they are strongly sensitized, indicating that additional pathogenic mechanism(s) operate in conjunction with Th2 immunity to drive asthma pathogenesis.

**Objectives:** Employ systems level analyses to identify inflammation-associated gene networks operative at baseline in sputum-derived RNA from house dust mite-sensitized (HDM^s^) subjects with/without wheezing history; identify networks characteristic of the ongoing asthmatic state. All subjects resided in the constitutively-HDMhigh Perth environment.

**Methods:** Genome wide expression profiling by RNASeq followed by gene coexpression network analysis.

**Measurements/Results:** HDM^s^-nonwheezers displayed baseline gene expression in sputum including IL-5, IL-13 and CCL17. HDM^s^-wheezers showed equivalent expression of these classical Th2-effector genes but their overall baseline sputum signatures were more complex, comprising hundreds of Th2-associated and epithelial-associated genes, networked into two separate coexpression modules. The first module was connected by the hubs EGFR, ERBB2, CDH1 and IL-13. The second module was associated with CDHR3, and contained genes that control mucociliary clearance.

**Conclusions:** Our findings provide new insight into the inflammatory mechanisms operative at baseline in the airway mucosal microenvironment in atopic asthmatics undergoing natural perennial aeroallergen exposure. The molecular mechanism(s) that determine susceptibility to asthma amongst these subjects involve interactions between Th2-and epithelial function-associated genes within a complex co-expression network, which is not operative in equivalently sensitized/exposed atopic non-asthmatics.

**Funding:** This study was funded by the Asthma Foundation WA, the Department of Health WA, and the NHMRC. AB is funded by a BrightSpark Foundation McCusker Fellowship. GLH is a NHMRC Fellow. AG is supported by the McCusker Charitable Foundation Bioinformatics Centre. ACJ is a recipient of an Australian Postgraduate Award and a Top-Up Award from the University of Western Australia.

## INTRODUCTION

Asthma is a chronic disease of the conducting airways that is characterized by airways inflammation, airways remodeling, and progressive loss of lung function. It is increasingly recognized as a highly heterogeneous disorder comprising multiple sub-phenotypes (1). The atopic form of the disease develops in early childhood, and is initiated by sensitization to inhalant allergens exemplified by house dust mite (HDM). Progression of atopic asthma towards chronicity is driven by repeated cycles of airways inflammation, in particular severe exacerbations triggered by respiratory infections which involve interactions between host anti-viral and atopy-associated effector mechanisms (2, 3), and the rate of the ensuing decline in lung function is related to the frequency and intensity of these exacerbations (4- 6).

Recent clinical intervention studies, including those demonstrating that treatment with anti-IgE reduces exacerbation frequency, confirms the causal role of Th2 responses in these intermittent events (7–9). However the degree to which chronic exposure to Th2-stimulatory perennial aeroallergens contributes to the inflammatory milieu in the airway mucosa of sensitized atopics during the periods between overt exacerbation events, thus potentially influencing long-term persistence of the asthma-associated wheezy phenotype, remains unclear. This is an important issue in relation to design of future therapeutic strategies for prevention of asthma progression i.e. is it sufficient to target severe exacerbation events alone, or is it potentially necessary to also dampen ongoing aeroallergen-driven Th2 reactivity at baseline in sensitized/perennially exposed subjects?

We have addressed this issue in a study population consisting of 22yr olds from an unselected birth cohort resident in Perth Western Australia (10). We have previously shown that the dominant asthma-associated aeroallergen in this region is HDM (11) which is present in local households at high levels throughout the year (12), and accordingly the study focused primarily on atopics who were sensitized and chronically exposed to HDM. Our approach was based on the recent demonstration that induced sputum, which contains a sample of cell populations present on the airway surface, can potentially be used for gene expression profiling of wheeze-associated inflammatory responses in asthmatics (13, 14).

In the present investigation we have employed RNA-Seq in conjunction with coexpression network analysis to profile asthma-associated gene networks in sputum samples collected at (symptom-free) baseline from study groups matched for age, HDM sensitization status and environmental exposure, but dichotomous with respect to wheezing symptom expression. Our findings suggest that upregulation of Th2 signature genes exemplified by the effector cytokines IL-5 and IL-13 is a common feature across the whole HDM^s^/exposed population at baseline, but in the subgroup with history of current wheeze the Th2 signature is more complex and intense, and is uniquely networked with a series of concomitantly upregulated epithelial cell associated pathways.

## METHODS

### Study population

This study was conducted within the 22-year follow-up of an unselected longitudinal birth cohort recruited in Perth, Western Australia, namely the Western Australia Pregnancy Cohort (Raine study, (10, 11)). The 22-year follow-up included 1234 active participants. Subjects were selected for case/control studies based on their clinical characteristics and the availability of high quality sputum samples (see below). Four clinical groups were defined; (i) HDM sensitized atopics (SPT ≥ 3.0mm) with current wheeze during previous 12mths, with or without a physician diagnosis of *“*asthma ever*”* (HDM^s^ wheezers, n=16); (ii) HDM sensitized atopics without current asthma or wheeze (HDM^s^ nonwheezers, n=24); (iii) nonatopics with current asthma and/or wheeze (nonatopic wheezers, n=7); (iv) nonatopics without current asthma or wheeze (nonatopic controls, n=21).

### Sputum induction and processing

Induced sputum was obtained after mannitol inhalation challenge (15). The samples were stored at 4 °C for up to 2 hours prior to processing. Sputum was processed (see the online data supplement) by selection and subsequent disruption of mucus plugs with forceps to minimize contamination with saliva (16).

### Transcriptome profiling by RNA-Seq

Total RNA was extracted from good quality sputum (cell viability > 48%, squamous < 32%) employing TRIzol (Ambion) followed by RNeasy MinElute (QIAgen). The mean±sd RNA integrity number was 7.6 ±1.0 as assessed on the bioanalzyer (Agilent). RNA samples were shipped on dry ice to the Australian Genome Research Facility for library preparation (TruSeq Stranded mRNA Library Prep Kit, Illumina) and sequencing (Illumina HiSeq2500, 50-bp single-end reads, v4 chemistry). The raw data are available at the NCBI Short Read Archive (accession; SRP057350).

### RNA-Seq data analysis

The quality of the RNA-Seq data was assessed with the Bioconductor package Rqc (see Fig. E1 in the online data supplement). Reads were aligned to the reference genome (hg19) using Subread, and summarized at the gene-level using featureCounts (17). Genes with less than 300 total counts were removed from the analysis. Differentially expressed genes were identified employing Empirical analysis of digital gene expression data in R (EdgeR) with False Discovery Rate (FDR) control for multiple testing (18). The analysis was adjusted for latent variation using the Remove Unwanted Variation (RUV) algorithm (Fig. E2 (19)). A coexpression network was constructed employing the weighted gene coexpression network analysis (WGCNA) algorithm (20). Prior to network analysis, the count data was transformed using the variance stabilizing transformation algorithm (18). Modules associated with clinical traits were identified by plotting the −log10 p-values from the edgeR analysis on a module-by-module basis. The wiring diagram of selected gene networks was reconstructed employing two different methods. The first method utilized “prior knowledge” comprising experimentally supported molecular relationships based on data from the Ingenuity Systems KnowledgeBase (http://www.ingenuity.com) (20). The second method utilized unbiased connectivity patterns derived from WGCNA, and the network was visualized using VisANT (21). Biological pathways and functions enriched in the data were identified with Enrichr (22).

### Immunostaining

Primary bronchial epithelial cells were obtained from 8 healthy nonatopic children and 8 atopic asthmatic children with HDM allergy who were undergoing elective surgery for non-respiratory related conditions. Cytospins were prepared and stained for CDHR3 and DAPI using methods previously described (see online supplementary methods).

## RESULTS

The characteristics of the 4 study groups are illustrated in Table 1

**Table 1.**
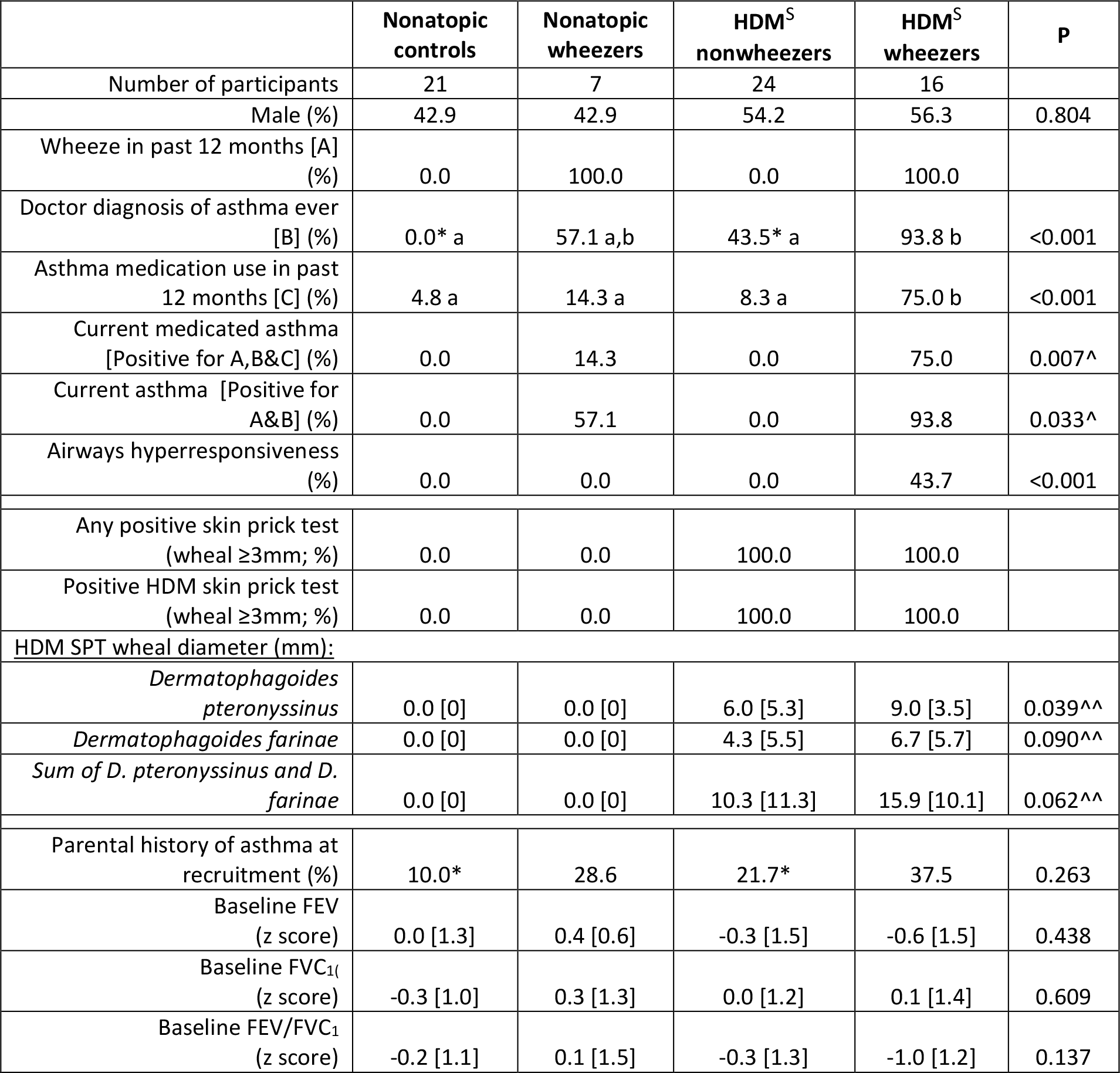
Characteristics of the study population.

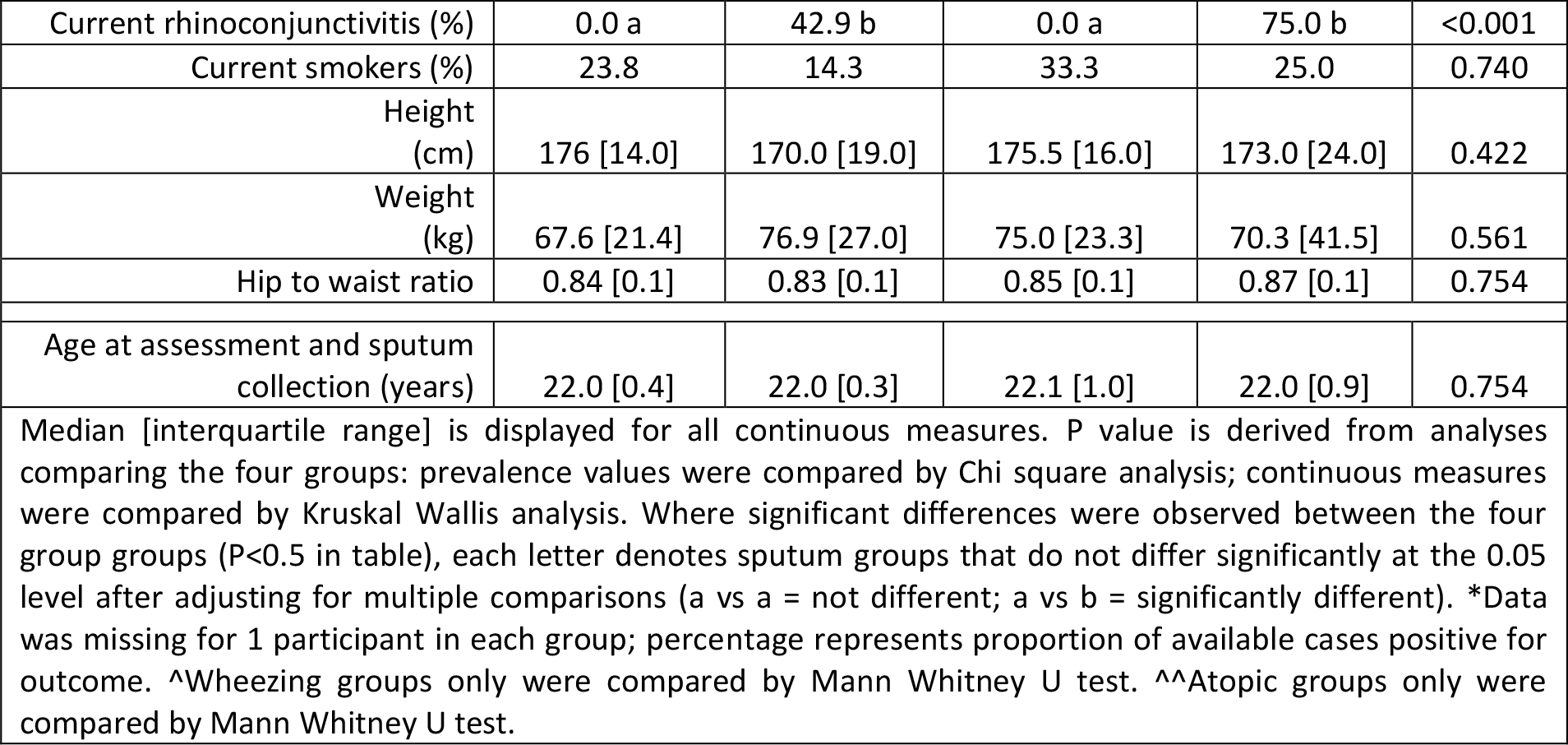

The cellular composition of sputum from these subjects was dominated by macrophages and neutrophils, which constituted 88-94% of the overall population, and the proportion of these cell types (and overall total yields) did not differ between the groups (see supplementary Table E1). Squamous cells and lymphocytes comprised on average 4.5% and 1.8% respectively and also did not differ between groups. Small numbers of eosinophils were detectable only in the atopic groups and were highest in the wheezers (Table E1).

Gene expression patterns in sputum were firstly compared between HDM^s^ nonwheezers and nonatopic controls. The data showed that 80 genes were upregulated (including the Th2 signature genes IL-5 [4.15 logfold] and IL-13 [2.92 logfold]) and 11 genes were downregulated (FDR < 0.05, Table E2). To obtain detailed information on the regulatory interactions between these genes, we utilized experimentally supported findings from published studies (prior knowledge) to reconstruct the underlying network (20). This analysis showed that the genes were mainly involved in IL-1B and IL-5/IL-13 signaling (Fig 1).Secondly, comparing gene expression patterns between HDM^s^ wheezers and nonatopic controls showed that 842 genes were upregulated (again including IL-5 [5.29 logfold], IL-13 [3.03 logfold] and IL-33 [2.59 logfold]) and 11 genes were downregulated in the wheezers (FDR < 0.05, Table E3). As illustrated in Fig. 2, the prior knowledge network revealed that these genes revolved around a few hubs-erb-b2 receptor tyrosine kinase 2 (ERBB2/HER2), which was involved in 59 interactions (also known as ‘edges’ in graph theory (23)), IL-13 (44 edges), and E-Cadherin/CDH1 (37 edges).

**Figure 1:**
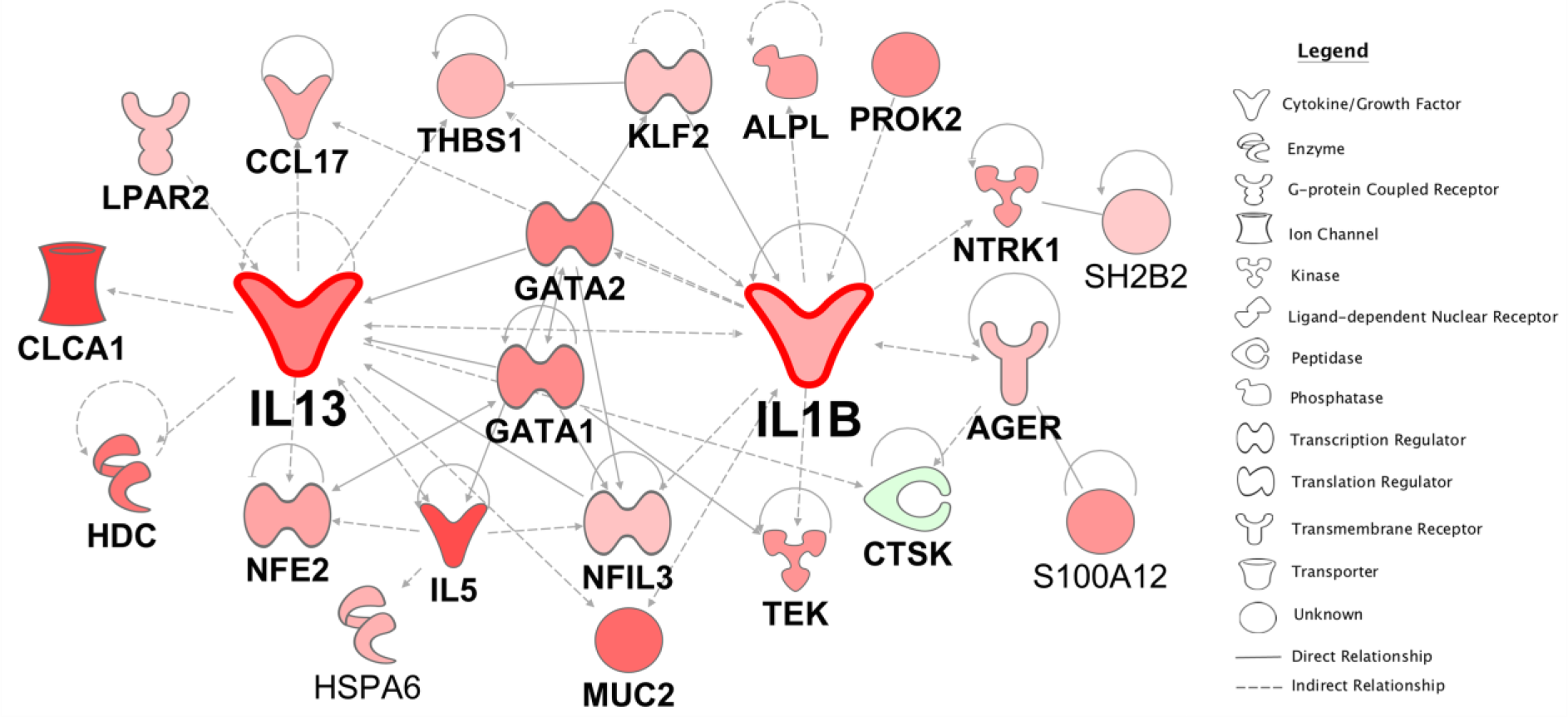
Gene expression patterns in sputum were compared between HDM^s^ nonwheezers and nonatopic controls. The network was reconstructed employing prior knowledge from the literature. Genes highlighted in red denote upregulation, whilst green indicates downregulation in HDM^s^ nonwheezers.

**Figure 2:**
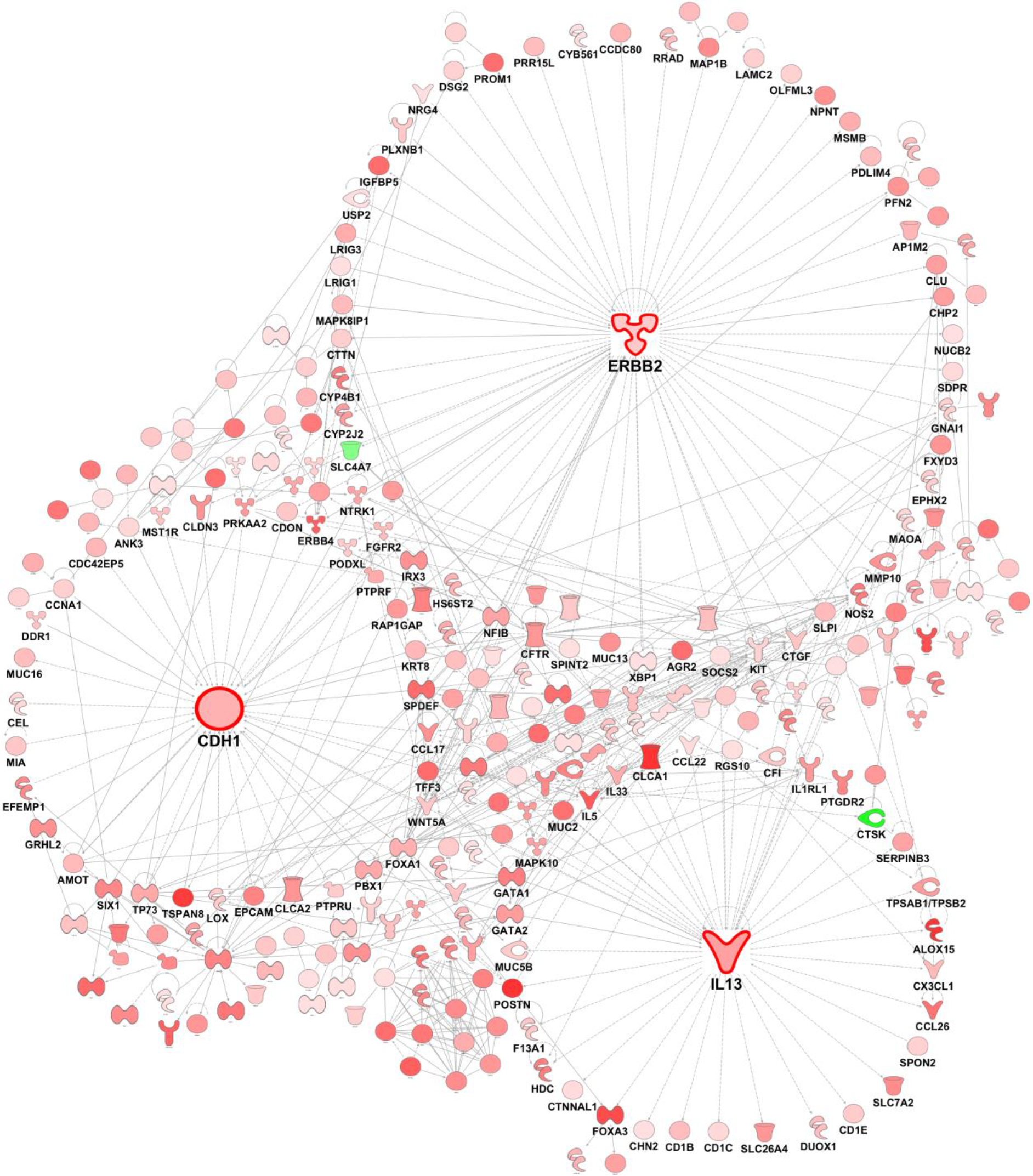
Gene expression patterns in sputum were compared between HDM^s^ wheezers and nonatopic controls. The gene network was reconstructed using prior knowledge. Genes highlighted in red denote upregulation, whilst molecules in green indicate downregulation in HDM^s^ wheezers.

Thirdly, we compared gene expression patterns between HDM^s^ wheezers versus HDM^s^ nonwheezers. The data showed that 859 genes were upregulated and 8 genes were downregulated (FDR < 0.05, Table E4). The prior knowledge network constructed from these genes (Fig. 3) identified epidermal growth factor receptor (EGFR, 60 edges), ERBB2 (56 edges) and CDH1 (38 edges) as hub genes. It is noteworthy that IL-13 did not feature here since it was not differentially expressed after adjustment for multiple testing (p-value = 0. 028, FDR = 0.27). In contrast, IL-33 was upregulated in this comparison (2.74 logfold; Table E4).

**Figure 3:**
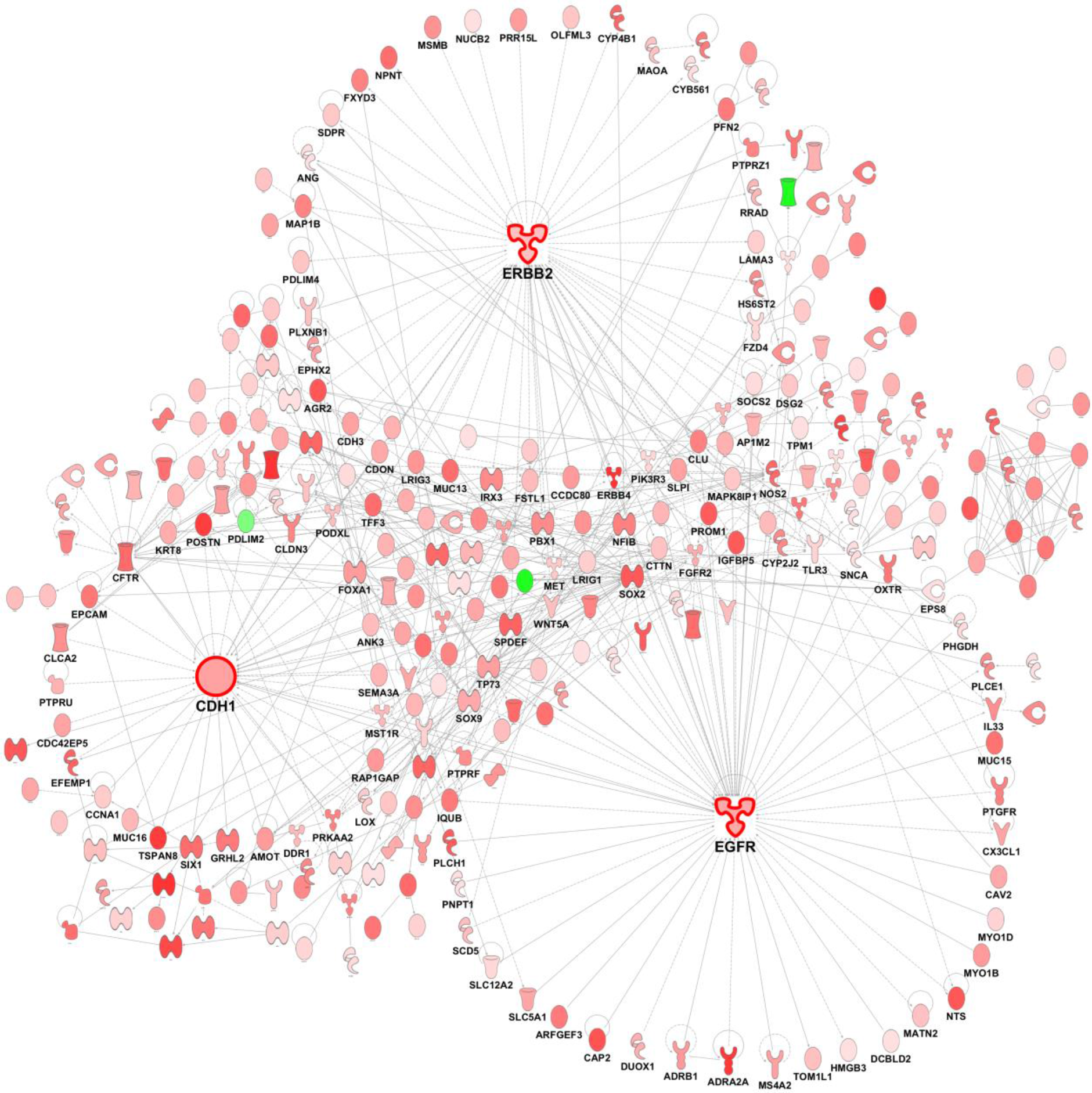
Gene expression patterns in sputum were compared between HDM^s^ wheezers versus HDM^s^ nonwheezers. The network was reconstructed employing prior knowledge from the literature. Genes highlighted in red denote upregulation, whilst molecules in green indicate downregulation in HDM^s^ wheezers.

Finally, we compared gene expression in nonatopic wheezers with nonatopic controls, and a single gene - LIM domain binding 3 (LDB3), was upregulated in the subjects with wheeze (FDR = 2.6 × 10^−5^). As expected, there was no evidence of a Th2 signature.

It has been reported that hub genes in biological interaction networks often exhibit limited expression changes in experimental asthma models (24), thus a potential caveat of the above analyses, which focused on differentially expressed genes, is that some hubs may have escaped detection. To address this issue, we constructed a genome-wide coexpression network, utilizing the data from both atopic groups (n=40). The resulting network comprised 14,833 genes organized into 23 coexpression modules. To identify disease-associated modules, we plotted the −Log10 p-values derived from the above differential expression analyses on a module-by-module basis. The data showed that the modules were not different between HDM^s^ nonwheezers and nonatopic controls (Fig E5A). In contrast, three modules (designated A, P, and Q) were upregulated in HDM^s^ wheezers versus the other two groups (nonatopic controls, HDM^s^ nonwheezers, Fig E5B, E5C).

Module “P” contained 319 genes, and the prior knowledge network constructed from these genes contained the hub genes EGFR (35 edges) and CDH1 (31 edges, Fig E6). Module “Q” contained 440 genes, and the hubs in the resultant prior knowledge network were ERBB2 (35 edges) and IL-13 (27 edges, Fig E7). Principal component analysis showed that these two modules (P, Q) were highly correlated (Pearson correlation: 0.897, P-value = 4.441 × 10^−15^) (Fig E8), suggesting they are subunits of a larger parent module. Therefore, we merged them into a single network. In the merged network the dominant hubs were EGFR (73 edges), ERBB2 (65 edges), CDH1 (56 edges) and IL-13 (48 edges, Fig E9). Notably, these hubs connect to both common and unique pathways (Fig 4). The biological function of the genes that interact with the hubs was interrogated using Gene Ontology terms (Table E5) and Pubmed searches (Table E6). Module “A” contained 506 genes. It was not possible to reconstruct this module using prior knowledge, because no interactions were found for the vast majority of genes. Therefore we used unbiased correlation patterns to reconstruct the network (21). This analysis showed that the highest-ranking coexpression hubs were TEKT1, FOXJ1, ARMC3, PIFO, DNAH5, RSPH1, FAM81B, SNTN, CDHR3, ERICH3, DNAH9, and CAPSL (Fig 5). This module was strongly enriched with genes involved in the function of ciliated epithelial cells (Table E7).

**Figure 4:**
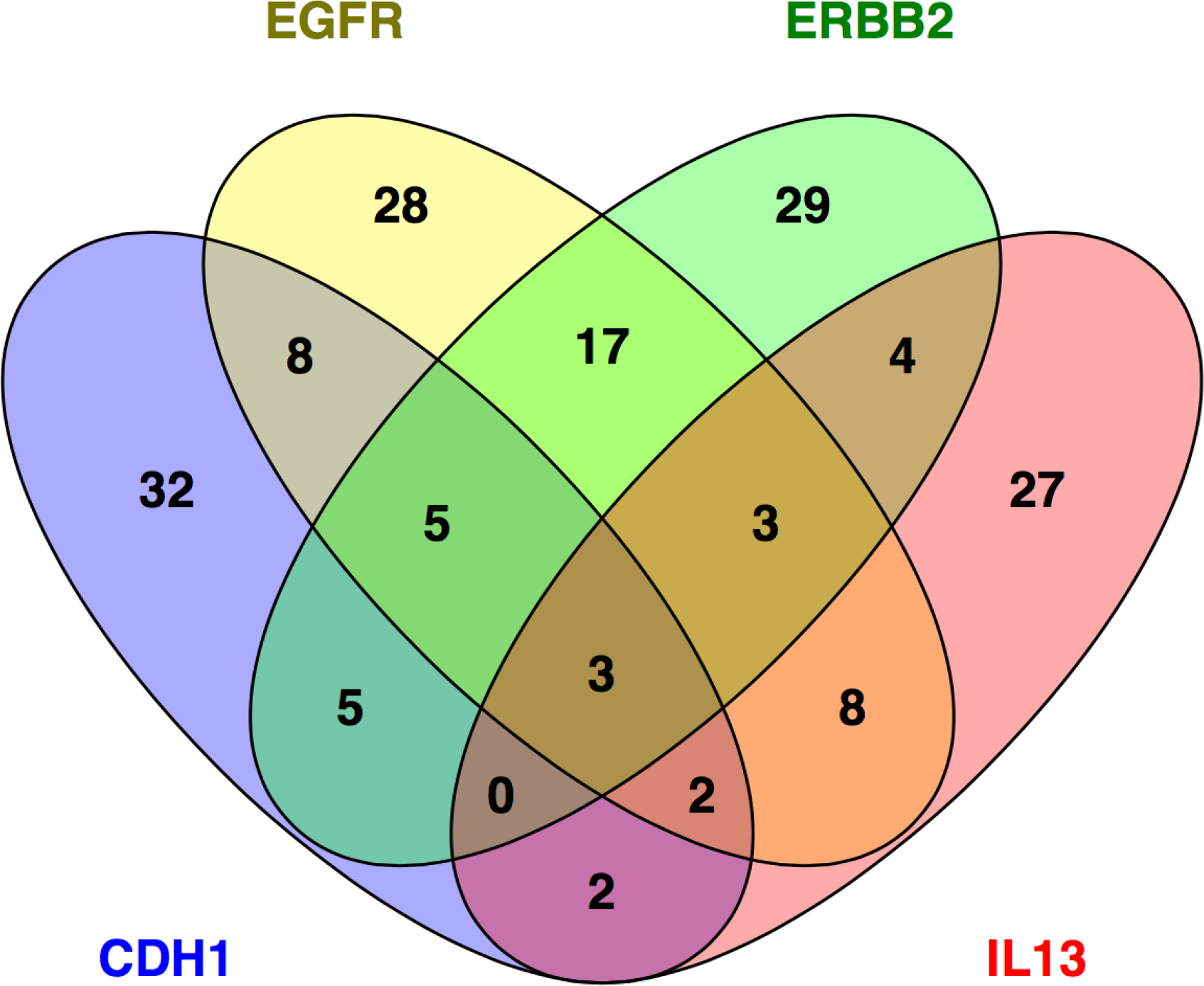
The Venn diagram illustrates the overlap between the genes that are networked with each hub.

**Figure 5:**
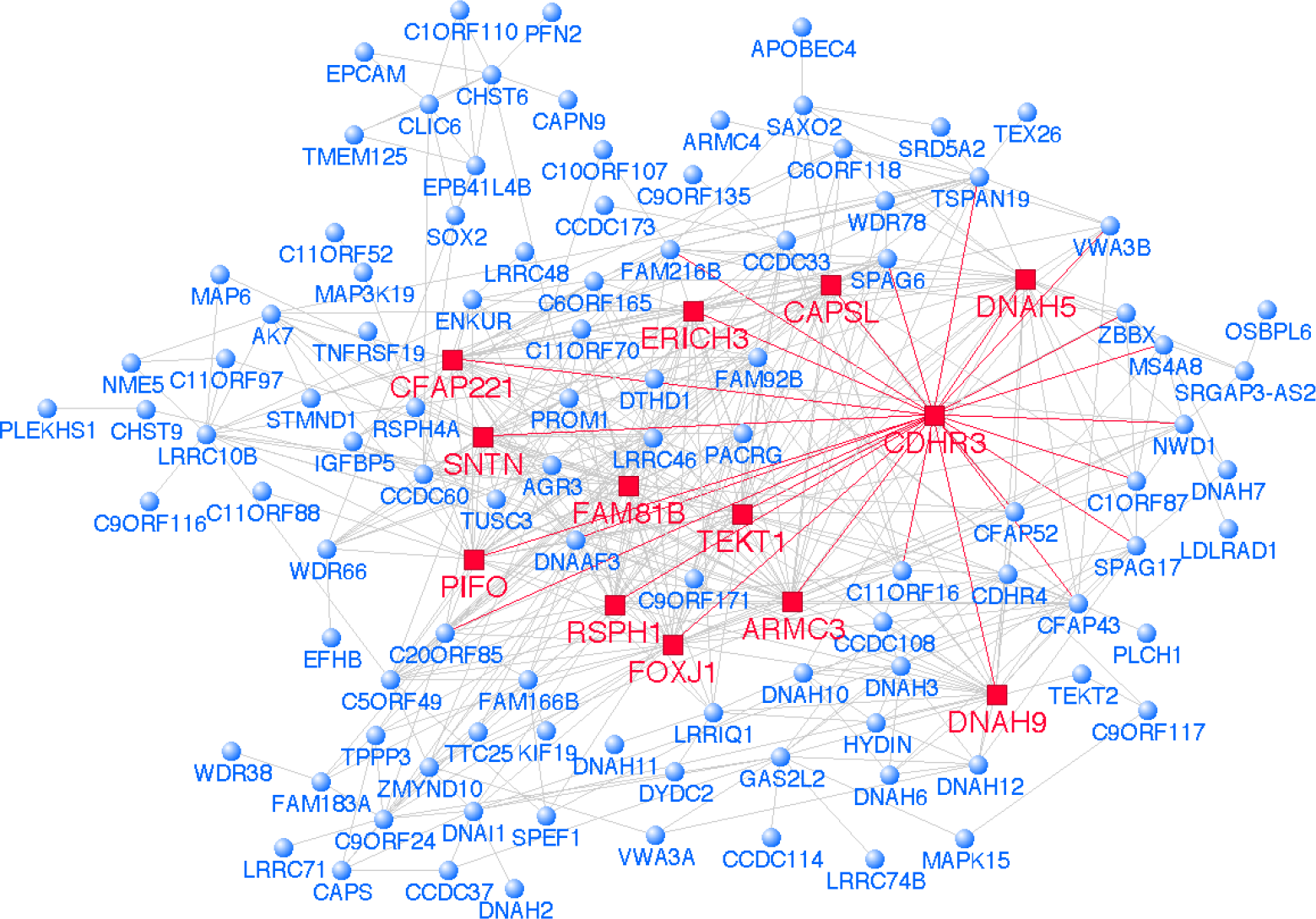
Reconstruction of the mucociliary clearance module identifies CDHR3 as a hub. This module was reconstructed with weighted correlation network analysis (WGCNA). The dominant hubs are highlighted in red.

We selected CDHR3 for further study because our data suggests it is a hub that functions in ciliated epithelial cells, and a previous study reported it was a susceptibility gene for severe asthma exacerbations (25). CDHR3 expression was examined in bronchial epithelial cells from HDM sensitized children with asthma (n=8) and from nonatopic controls (n=8) using immunostaining (see Table E7 for subject characteristics). The data showed there was positive staining localized to the apical surface of columnar epithelial cells in both cohorts (Fig. 6A, Fig E10). Of particular interest was the observation that expression of CDHR3 (green) appeared more intense and defined in airway epithelial cells derived from the asthmatic children, and this was confirmed by image quantification (Fig. 6B).

**Figure 6:**
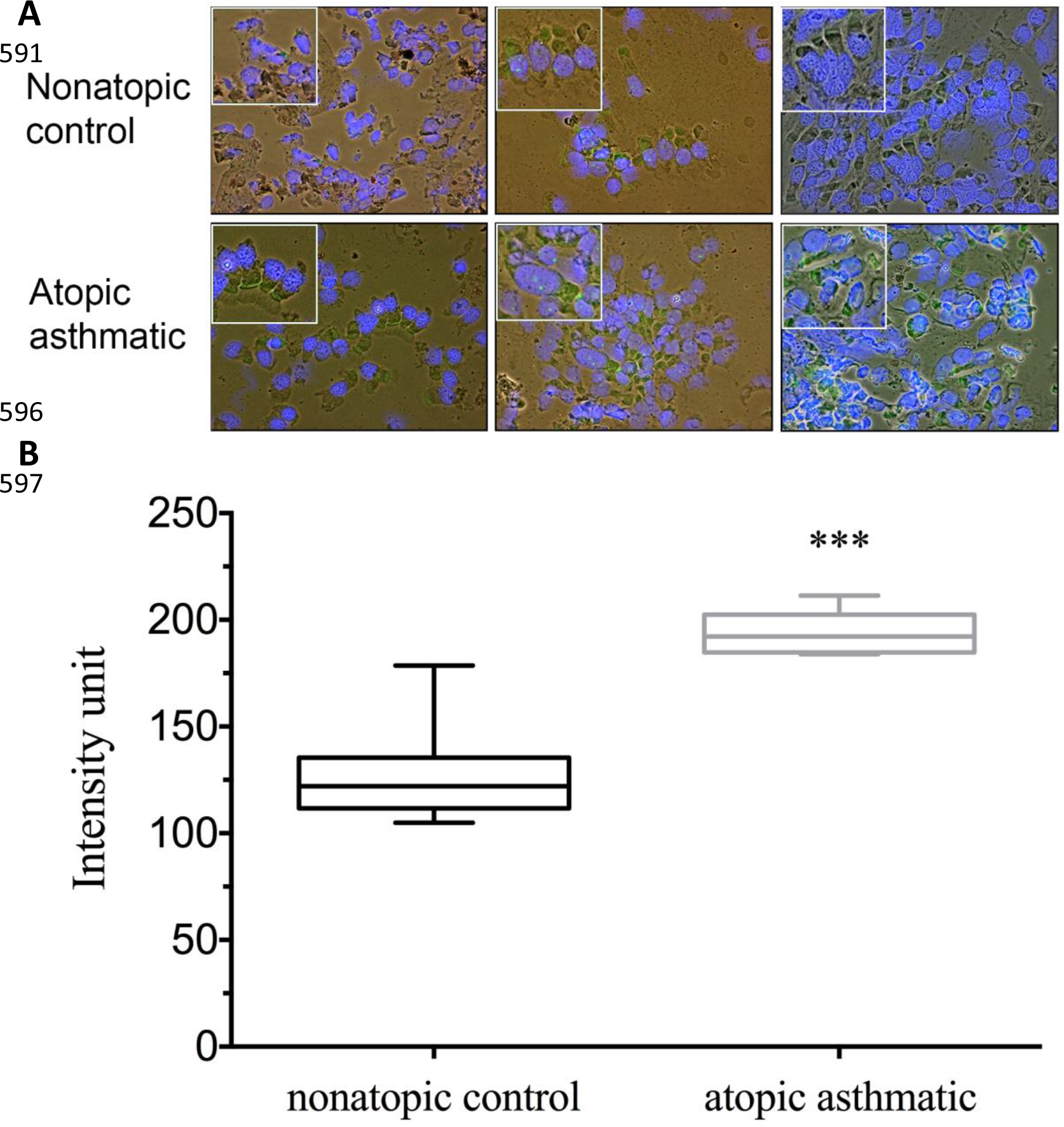
Expression of CDHR3 in bronchial epithelial cells from HDM sensitized atopics with asthma and nonatopic controls. A) Bronchial epithelial cells were immunofluorescently stained for CDHR3 expression (green) and nuclei with DAPI (blue). Staining images were then overlaid over bright field images taken of the same field of view. Note: mag 200x; inset 400x. B) Quantification of the images demonstrated that the expression was more intense in the atopics with asthma. *** P-value < 0.001

## DISCUSSION

An increasing body of epidemiological and experimental evidence (reviewed (2, 3, 26)), now supported by a range of intervention studies (4–6), argues for a causal role for Th2-associated inflammatory mechanisms in the aetiology and pathogenesis of atopic asthma. However the precise details of the underlying causal pathways still remain incompletely understood. In particular, the relative contributions of airways inflammation resulting from acute severe exacerbation events versus chronic exposure to relevant aeroallergens to time-related lung function decline in asthmatics, remains unknown. Moreover, while it is undisputed that sensitization to perennial aeroallergens is an important asthma risk factor, community wide studies clearly demonstrate that only a minority of sensitized subjects (including of those highly sensitized to HDM (11)) ever develop persistent wheeze. This suggests that additional cofactor(s) may be required to unmask the full pathogenic potential of aeroallergen-specific sensitization. A likely candidate in this regard is the airway epithelium which may function as both a target for Th2-associated inflammation and/or as an active participant via production of a range of immunomodulatory molecules that can regulate the local functioning of Th2 cells and also Th2 cytokine-secreting group 2 innate lymphoid cells (7–9).

Our current study design represents an unbiased approach towards testing this possibility. In the core experiments we have sampled induced sputum cell populations from equivalently sensitized adult atopics undergoing natural aeroallergen exposure, and subsequent gene expression profiling and ensuing bioinformatics analyses after stratification on the basis of wheezing phenotypes provides novel insight into the nature of the inflammatory processes ongoing on the airway mucosal surface at the time of sampling.

Our initial analyses showed that a Th2 gene expression program was upregulated in baseline sputum samples from the HDM sensitized atopics, regardless of whether these subjects have current history of wheeze. The key Th2 genes IL-5 and IL-13 were upregulated to comparable degrees in both groups, however in HDM^s^ nonwheezers the overall Th2 program was restricted to only a small number of IL-5/IL-13-associated genes. In contrast, hundreds of genes were upregulated in HDM^s^ wheezers and network analysis suggested that these genes function in the context of two discrete coexpression modules. Reconstruction of the first module using prior knowledge revealed that the hub genes EGFR, ERBB2, CDH1, and IL-13 dominated the network structure. The second coexpression module comprised genes that control mucociliary clearance, and reconstruction of this module employing unbiased gene coexpression patterns identified CDHR3 as a hub. Overall, our findings suggest that the molecular mechanisms that determine susceptibility to asthma-associated wheeze amongst HDM sensitized atopics involve complex interactions between Th2 and epithelial gene networks.

EGFR was the dominant hub in the first module. Downstream of this gene is a complex signaling pathway that can be activated by multiple ligands (e.g. amphiregulin, EGF, epiregulin, HB-EGF, TGF-a) (27). Puddicombe et al. reported that EGFR was upregulated in the bronchial epithelium of patients with asthma and in particular severe asthma in comparison to healthy controls, and expression levels were correlated with sub-epithelial reticular membrane thickening (28). Le Cras et al. reported that inhibition of EGFR signaling with a tyrosine kinase inhibitor reduced goblet cell hyperplasia, airway hyperreactivity and airway smooth muscle thickening in a chronic mouse model of HDM exposure (29). The latter two phenotypes were also reduced by conditional transgenic expression of a dominant negative EGFR mutant in the lung epithelium. Together, these data suggest that upregulation of EGFR signaling networks in the context of HDM exposure plays a causal role in the development of asthma-related traits.

The second hub ERBB2 is an orphan receptor from the EGFR family. It lacks a ligand-binding domain and transduces signals by forming heterodimers with other ligand bound members of the EGF receptor family, including EGFR. Polosa et al. reported that ERBB2 expression was not different in bronchial epithelial cells from asthmatic subjects compared to healthy controls (30). Song and Lee identified ERBB2 as an asthma susceptibility gene based on a pathways analysis of genome-wide single nucleotide polymorphism data (31). The function of ERBB2 in asthma has not been previously investigated in animal models. Vermeer et al. reported that blockade of ERBB2 signaling in differentiated airway epithelial cells cultured at air-liquid interface reduced the number of ciliated epithelial cells (32). Kettle et al. reported that blocking ERBB2 signaling *in vitro* attenuated neuregulin-induced upregulation of MUC5AC and MUC5B (33). Notably, our network analysis showed that ERBB2 connects to anterior gradient 2 (AGR2). Previous studies have shown that ERBB2 upregulates the transcription and secretion of AGR2 (34, 35). AGR2 binds to immature MUC5AC in the endoplasmic reticulum, where it is thought to play a role in mucin folding. AGR2 deficient mice have profound defects in intestinal mucus production and reduced mucus production in the airways of allergen challenged mice (36, 37). Upregulation of ERRB2 networks may therefore influence asthma by modulating epithelial differentiation and mucus production.

The third hub E-cadherin (CDH1) is a cell adhesion molecule that forms adherence junctions between adjacent airway epithelial cells and maintains epithelial barrier integrity (38). HDM disrupts epithelial barrier function by delocalizing E-cadherin and other junction molecules, and this is thought to enhance allergic sensitization and inflammation (39). Polymorphisms in CDH1 are associated with airways remodeling and lung function decline, but only in those asthma patients using corticosteroids (40). Dysregulation of CDH1 signaling networks may impact on barrier function, inflammation, and airways remodeling.

The fourth hub IL-13 plays a central role in the pathogenesis of asthma by driving mucus production, airways hyper-responsiveness, and airways remodeling (26). It is produced by Th2 and group 2 innate lymphoid cells (ILC2), and it can also be produced by macrophages (41, 42). IL-13 itself was not differentially expressed in HDM^s^ wheezers versus nonwheezers, however network analysis demonstrated that in the wheezers it was connected to an extensive set of genes that have established roles in mouse models of allergic asthma. For instance, IL-33 stimulates the production of IL-5 and IL-13 by type 2 innate lymphoid cells and Th2 cells (43, 44), and in the presence of GM-CSF it can drive allergic inflammation at sub-threshold allergen doses (45). In animal models, deficiency of multiple genes from the IL-13 network can impact on asthma-related traits, including allergic sensitization and/or inflammation (ALOX15 (46), CYBB (47)), and airways hyperresponsiveness and mucus production/goblet cell hyperplasia (POSTN (48), SERPINB3/4 (49)). Moreover, transgenic expression of SPDEF or FOXA3 leads to upregulation of pulmonary Th2 cytokines and increased goblet cell differentiation, eosinophilic inflammation, and airway hyperresponsiveness (50). It is noteworthy, that whilst both IL-13 and EGFR ligands can induce the transcription of mucin genes, microarray profiling studies have shown that these pathways have largely independent effects on gene regulation in bronchial epithelial cells, and they play distinct roles in goblet cell metaplasia (36, 51, 52). Many other pathways were also identified that are regulated by IL-13 and relevant to asthma pathogenesis (e.g. CCL17, CCL26, CTGF, FCER1A, KITLG, MUC2, NOS2, TLR3, see Table E5).

The second coexpression module we identified comprised genes expressed in ciliated epithelial cells that control mucociliary clearance. The primary function of cilia is to beat in a synchronous manner to clear mucus from the airways and into the pharynx. Thomas et al. reported that cilia beat frequency was decreased in patients with asthma, and severe asthmatics had abnormal ciliary orientation and microtubule defects (53). Notably, employing network analysis we showed that CDHR3 was a highly ranked coexpression hub within this module. This prompted us to examine CDHR3 protein expression in bronchial epithelial cells, and we demonstrated that expression was localized to the apical surface of columnar epithelial cells and was increased in HDM sensitized atopics with asthma compared to nonatopic controls. Ross et al. reported that CDHR3 was highly upregulated during mucociliary differentiation of human airway epithelial cells (54). Bisgaard and coworkers reported that polymorphisms in CDHR3 were associated with recurrent, severe childhood asthma exacerbations (25). More research will be required to investigate the role of CDHR3 in ciliated epithelial cells.

This exploratory study has limitations including small sample size that should be acknowledged. The molecular profiling studies were based on a heterogeneous cell population, and the pathways we identified were mainly associated with airway epithelial cells, which represent a minority population in sputum. We cannot exclude the possibility that epithelial shedding may have varied across the study groups and impacted on the analysis, although the RUV adjustment we employed should minimize any potential confounding by biological and/or technical variations. It is additionally noteworthy that using prior knowledge to reconstruct gene networks relies on data derived from experimental settings that may be far removed from the current study, which means that conclusions drawn from these analyses may be oversimplified given that genes can function in a context specific manner. Detailed follow-up mechanistic will therefore be required to elucidate the specific cellular mechanisms involved, and dissect the role of the molecular pathways we have identified.

Notwithstanding these caveats, our findings collectively are consistent with the general hypothesis that progression from subclinical responsiveness to aeroallergen exposure in atopic asthmatics to expression of the persistent wheezing phenotype involves the establishment of coexpression networks linking Th2 effector cytokine genes in immune cells recruited to the airway surface with genes expressed in adjacent epithelial cells that have been implicated in myriad asthma-relevant functions including mucosal barrier integrity, mucus production, tissue remodeling, responsiveness to irritants, and (exemplified by IL-33) intensification of aeroallergen-specific Th2 immunity. Targeting drug development programs specifically at these chronic mechanisms, as opposed to simply those that are triggered during acute exacerbation events, may provide improved therapeutics for prevention of asthma progression in atopics who represent the segment of the population at greatest risk of this disease.

## Acknowledgements

We would like to acknowledge the participants of the Raine Study for their ongoing participation in the study, the Raine Study Team for co-ordination of the study and data collection, and the UWA Centre for Science and the Sleep Study Technicians. We would like to acknowledge the University of Western Australia (UWA), Curtin University, the Raine Medical Research Foundation, the UWA Faculty of Medicine, Dentistry and Health Sciences, the Telethon Kids Institute, the Women’s and Infant’s Research Foundation (King Edward Memorial Hospital) and Edith Cowan University for providing funding for the Core Management of the Raine Study. The 22-year Raine Study follow-up was funded by NHMRC project grants 1027449, 1044840 and 1021858. Funding was also generously provided by Safework Australia. We would like to acknowledge the funding support of Pharmaxis, Australia, for the provision of the Aridol challenge kits used in this study. Pharmaxis did not have any input into the study design, data collection, interpretation and preparation of this manuscript.

